# A rapid and accurate MinION-based workflow for tracking species biodiversity in the field

**DOI:** 10.1101/617019

**Authors:** Simone Maestri, Emanuela Cosentino, Marta Paterno, Hendrik Freitag, Jhoana M. Garces, Luca Marcolungo, Massimiliano Alfano, Iva Njunjić, Menno Schilthuizen, Ferry Slik, Michele Menegon, Marzia Rossato, Massimo Delledonne

**Author notes:** S.M. and E.C. contributed equally.

## Abstract

Genetic markers (DNA barcodes) are often used to support and confirm species identification. Barcode sequences can be generated in the field using portable systems based on the Oxford Nanopore Technologies (ONT) MinION platform. However, to achieve a broader application, current proof-of-principle workflows for on-site barcoding analysis must be standardized to ensure reliable and robust performance under suboptimal field conditions without increasing costs. Here we demonstrate the implementation of a new on-site workflow for DNA extraction, PCR-based barcoding and the generation of consensus sequences. The portable laboratory features inexpensive instruments that can be carried as hand luggage and uses standard molecular biology protocols and reagents that tolerate adverse environmental conditions. Barcodes are sequenced using MinION technology and analyzed with ONTrack, an original *de novo* assembly pipeline that requires as few as 500 reads per sample. ONTrack-derived consensus barcodes have high accuracy, ranging from 99,8% to 100%, despite the presence of homopolymer runs. The ONTrack pipeline has a user-friendly interface and returns consensus sequences in minutes. The remarkable accuracy and low computational demand of the ONTrack pipeline, together with the inexpensive equipment and simple protocols, make the proposed workflow particularly suitable for tracking species under field conditions.

## 1. Introduction

Recent advances in molecular biology allow the use of genetic markers (DNA barcodes) to support and confirm morphological evidence for species identification and to quantify interspecific differences in order to compare species in terms of evolutionary distance. Most barcodes are still generated using the Sanger sequencing method, which requires access to a well-equipped molecular biology laboratory. Second-generation sequencing technologies are also used for barcoding, but they depend on expensive equipment and the reads are often too short to distinguish species reliably. The third-generation sequencer Oxford Nanopore Technologies (ONT) MinION based on nanopores has proven successful for sequencing under extreme field conditions such as the tropical rainforests of Tanzania, Ecuador and Brazil [1–3], the hot savannah of West Africa [4], and the ice floes of Antarctica [5]. Bringing the laboratory to the field avoids the transport of samples to sequencing facilities, thus greatly reducing the analysis time and the need to export genetic material from collection sites.

Although several groups have reported successful on-site barcoding, it remains difficult to perform molecular biology procedures in sub-optimal and extreme environments. In our first expeditions, the quality of sequences generated in the field was consistently lower than achieved in the laboratory, suggesting that reagents and flow cells were affected by the unstable shipping and/or environmental conditions [1]. Furthermore, a recent on-site MinION run produced a low output consisting primarily of adapter sequences, probably reflecting the deterioration of the ligation enzyme and flow cells during suboptimal storage [2]. Some groups used lyophilized reagents to overcome adverse environments [1]. However, also equipment can be affected by extreme conditions, as we found on two different expeditions to Borneo during which one of the two models of portable PCR machine we brought with us lost temperature calibration resulting in the overheating and consequent failure in barcode amplification. The identification of robust protocols and equipment that tolerates suboptimal transport and operating conditions (but remains simple, inexpensive and portable) is therefore highly desirable in order to exploit the full potential of barcode sequencing in the field.

MinION-based sequencing is advantageous because it is portable, but it has a higher error rate than other methods and thus appropriate analysis workflows are therefore needed to generate high-quality barcode sequences [1,6]. High accuracy is particularly important in DNA-based taxonomy, as the threshold for intra-versus interspecific divergence of the COI gene is usually at about 2% [7] and in evolutionary ‘young’ species even lower [8]. We have previously attempted to reduce the high error rate of MinION by using more accurate 2D reads derived from the consensus of the forward and reverse strands. However, 2D sequencing kits are no longer available and have been replaced by 1D^2^ kits, which have yet to be optimized for amplicon sequencing. Even so, new ONT chemistries and software updates have greatly improved the throughput and 1D-read accuracy of nanopore sequencing in the last 2 years [8, 9]. Based on this reduced error rate (10–15%, R9.4 chemistry), several groups developed their own data analysis pipelines for barcoding, but none of the methods has yet achieved the status of ‘the gold standard’ [1,2,6,9].

Two main strategies are used to generate high-quality barcode sequences: reference-based and *de novo* pipelines. During the early development of nanopore sequencing, the high error rate in homopolymer runs made reference-based methods the better approach [1,2]. In a typical workflow, sequence reads are mapped to a reference sequence selected according to *a priori* knowledge, and the consensus sequence is ultimately determined based on the majority rule. Reference-based pipelines are useful when matching a target sequence to similar existing ones, but they struggle to reconstruct an accurate barcode if the organism of interest has not been sequenced before. Notably, if the target species carries an insertion compared to the reference species, the additional nucleotides are not included in the final consensus sequence [2]. Unlike the reference-based approach, *de novo* assembly pipelines rely only on the newly-generated reads. Therefore, they suffer more sequencing errors, especially if they are distributed in a nonrandom manner, and *ad hoc* error correction methods are needed to generate the barcodes using *de novo* assembly [2].

Recently, hybrid methods incorporating aspects of both approaches have been described [1,6]. One example is our *ONtoBAR* pipeline [1]. This creates a draft consensus sequence by assembling MinION reads *de novo* and uses the draft to retrieve the most similar sequence from the NCBI nt database, allowing the final consensus to be generated. Given the assumption that closely-related species differ mainly due to the accumulation of single-nucleotide polymorphisms (SNPs) rather than insertion/deletion polymorphisms (INDELs) that can generate frameshifts, the pipeline uses the reference sequence as a scaffold, allowing the correction of mismatches derived from MinION errors. Another hybrid method known as the *aacorrection* pipeline [6] is based on similar principles, in that a draft consensus sequence is used to recover matching sequences from the NCBI nt database. These are used to determine the correct reading frame, and generic bases (N) are introduced into the MinION-derived consensus in order to preserve amino acid assignments. A recent study compared reference-based and *de novo* approaches, finding that the *de novo* approach was more accurate because the reference-based approach can introduce bias by missing INDELs [2]. However, the filtering step in the proposed pipeline relied on quality scores (Q-scores) that are often recalibrated after basecaller updates, making the results strongly dependent on the sequencing chemistry and the basecaller version.

To fully exploit the potential of barcoding in the field, the proof-of-principle workflows reported thus far must be translated into standardized systems allowing on-site sequencing by professional users. Our involvement in conservation projects has motivated us not only to continuously improve the analytical precision of the pipeline in order to track biodiversity at the species level more accurately, but also to identify simple, rapid and inexpensive protocols. Here we demonstrate the results achieved using an updated barcoding workflow that features improvements both to the molecular biology field laboratory components and the subsequent data analysis.

## 2. Materials and Methods

### 2.1 Portable genomics laboratory

The portable genomics laboratory included the following equipment: three micropipettes (P1000, P200 and P20, Eppendorf), a mini-microcentrifuge (Labnet Prism Mini Centrifuge, Labnet), a thermal cycler (MiniOne PCR System, MiniOne), an electrophoresis system (MiniOne Electrophoresis System, MiniOne), a fluorometer (Qubit 2.0, Thermo Fisher Scientific), the nanopore sequencer (MinION, ONT) and an ASUS laptop (i7 processor, 16 GB RAM, 500 GB SSD) (Figure 1). The equipment was wrapped in air-bubble packaging, transported in a single Peli case (55×45×20 cm) (Figure 1) and checked as standard hold baggage in domestic and international flights (except the laptop, which was carried in the cabin). Standard molecular biology reagents were selected and used as described below. Reagents that required storage at 4 °C or –20 °C were transported in a foam box containing ice packs, and MinION flow cells were stored in a thermal bag in the same box. PCR primers were transported lyophilized and subsequently resuspended in 10 mM Tris-HCl (pH 8.0) supplemented with 1 mM EDTA and kept at room temperature.

**Figure 1.**
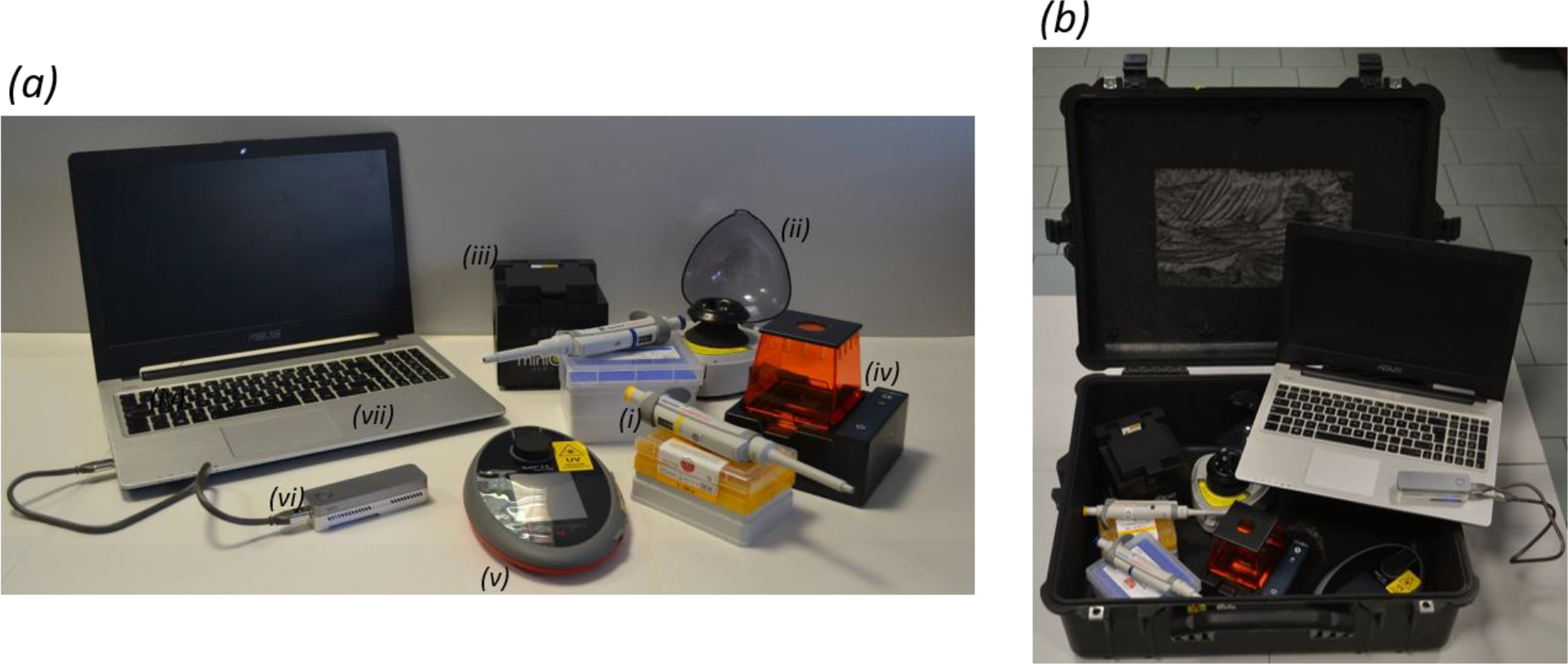
The portable genomics laboratory. Panel (a) shows the equipment comprising the portable genomics laboratory, namely (i) micropipettes, (ii) a mini-microcentrifuge, (iii) a thermal cycler, (iv) an electrophoresis system, (v) a fluorometer, (vi) the nanopore sequencer MinION, and (vii) a laptop. Panel (b) shows how the laboratory is transported.

### 2.2 Sample collection, DNA extraction and barcode amplification

Sample collection, tissue dissection, total DNA extraction, barcode amplification, MinION library preparation and sequencing were conducted in the field at the Ulu Temburong National Park (Brunei, Borneo) in October 2018, during a Taxon Expedition (https://taxonexpeditions.com/). We analyzed seven samples: two snails (Snail1 and Jap1) and five beetles (H36, H37, H42, H43 and Colen1). Two of them (H42, H43) were collected in an emergence trap [10] in which the specimens were exposed to a preserving agent consisting of ethanol (∼65%) glycerol (∼30%), water (∼5%) and a little amount of dish-washing detergent for several days.

Total genomic DNA was isolated using the DNeasy Blood and Tissue Kit (Qiagen) from a 1×1 mm biopsy of snail tissue or from the whole beetle after cutting the thorax and abdomen. Samples were incubated in ATL lysis buffer for 2 h at 56 °C and overnight at room temperature before DNA was extracted according to the manufacturer’s instructions and eluted in Tris-EDTA buffer (10 mM Tris, 1 mM EDTA, pH 8.0).

Barcoding PCR was conducted by amplifying the mitochondrial gene encoding cytochrome oxidase I (COI) using a MiniONE portable PCR device (MiniOne), lyophilized oligonucleotides and PCR reagents previously kept at room temperature. We used the universal primers LCO1490 and HC02198 [11] tailed with adaptors to allow indexing prior to MinION library preparation: 5’-TTT CTG TTG GTG CTG ATA TTG CGG TCA ACA AAT CAT AAA GAT ATT GG-3’ and 5’-ACT TGC CTG TCG CTC TAT CTT CTA AAC TTC AGG GTG ACC AAA AAA TCA-3’. Each PCR (total volume 25 μl) comprised 2 μl of the DNA template, 0.25 μM of each primer, 0.25 mM of each dNTP, 1× Herculase II reaction buffer, and 0.25 μl (20 U/μl) of Herculase II fusion DNA polymerase (Agilent Technologies). The amplification profile consisted of an initial denaturation step (3 min at 95 °C) followed by 35 cycles of 30 s at 95 °C, 30 s at 52 °C and 60 s at 72 °C, and a final extension for 5 min at 72 °C. PCR products were verified by electrophoretic analysis (MiniOne Electrophoresis System, MiniOne) for the presence of unique bands at the expected size (∼700bp). The amplification of H37 and Colen1 was not successful, so these samples were amplified using primers LepF1 (5’-TTT CTG TTG GTG CTG ATA TTG CAT TCA ACC AAT CAT AAA GAT ATT GG-3’) and LepR1 (5’-ACT TGC CTG TCG CTC TAT CTT CTA AAC TTC TGG ATG TCC AAA AAA TCA-3’) [12] using the reagents described above. The amplification profile consisted of an initial denaturation step (1 min at 95 °C) followed by six cycles of 1 min at 95 °C, 90 s at 45°C and 75 s at 72 °C, then 36 cycles of 1 min at 95 °C, 90 s at 51°C and 75 s at 72 °C and a final extension for 5 min at 72 °C. PCR products were purified using 1.5X AMPureXP beads (Beckman Coulter) and quantified using a Qubit 2.0 fluorimeter and the Qubit dsDNA BR assay kit (Thermo Fisher Scientific).

To incorporate index sequences and allow the sequencing of multiple samples in each MinION flow cell, a second round of PCR was carried out using 48 μl of the purified COI-PCR amplicons from the first round (0.5 nM), 2 μl of indexed primers provided in the EXP-PBC001 kit (ONT), 0.25 mM of each dNTP, 1× Herculase II reaction buffer, and 1 μl (20 U/μl) of Herculase II fusion DNA polymerase. The amplification profile consisted of an initial denaturation step (3 min at 95 °C) followed by 15 cycles of 15 s at 95 °C, 15 s at 62 °C and 30 s at 72 °C, and a final extension for 3 min at 72°C. Indexed PCR products were purified using 0.8X AMPureXP beads (Beckman Coulter), quantified as described above and pooled in equimolar concentrations.

### 2.3 MinION library preparation and sequencing

We used 1 μg of pooled amplicons to prepare sequencing libraries with the SQK-LSK108 DNA Sequencing kit (ONT) according to the manufacturer’s instructions (but omitting the DNA fragmentation step). The library was loaded on a FLO-MIN106 flow cell (R9.4 sequencing chemistry). Sequencing was carried out for 7 h in the field using MinKNOW v1.6.11 (ONT) on a portable laptop.

### 2.4 Sanger sequencing

Sanger sequencing was performed on COI PCR products prepared as described above and purified using 1X AMPureXP beads. Sequencing was carried out at the BMR Genomics facilities in Padova (Italy) or at the Museum für Naturkunde of Berlin (Germany), following our return from the field expedition. Forward and reverse Sanger reads were assembled into a consensus sequence using Geneious Prime v2019.0.4 (http://www.geneious.com/).

### 2.5 Bioinformatic analysis of MinION reads

After MinION sequencing, raw fast5 reads were basecalled and demultiplexed using Guppy v2.3.7+e041753. To reduce the number of misassignments, a second round of demultiplexing was performed requiring tags at both ends of reads using Porechop v0.2.3_seqan2.1.1 (https://github.com/rrwick/Porechop). Tags and adapters were trimmed using Porechop and reads of abnormal length were filtered out using a custom script.

Starting from pre-processed MinION reads, the *ONTrack* pipeline consisted of the following steps. First, VSEARCH v2.4.4_linux_x86_64 [13] was used to cluster reads at 70% identity and only reads in the most abundant cluster were retained for subsequent analysis. Next, 200 reads were randomly sampled using Seqtk sample v1.3-r106 (https://github.com/lh3/seqtk) and aligned using MAFFT v7.407 with parameters --localpair --maxiterate 1000, specific for iterative refinement, incorporating local pairwise alignment information [14]. EMBOSS cons v6.6.6.0 (http://emboss.open-bio.org/rel/dev/apps/cons.html) was then used to retrieve a draft consensus sequence starting from the MAFFT alignment. The EMBOSS cons plurality parameter was set to the value obtained by multiplying the number of aligned reads by 0.15, in order to include a base in the draft consensus sequence if at least 15% of the aligned reads carried that base. If less than 15% of the aligned reads carried the same base in a specific position, and a generic base (N) was included in the consensus sequence, the generic base was removed using a custom script. To polish the obtained consensus sequence, 200 reads were randomly sampled using Seqtk sample, with a different seed to the one used before, and mapped to the draft consensus sequence using Minimap2 v2.1.1-r341 [15]. The alignment file was filtered, sorted and compressed to the *bam* format using Samtools v1.7 [16]. Nanopolish v0.11.0 (https://github.com/jts/nanopolish) was used to obtain a polished consensus sequence. When the *ONTrack* pipeline was run multiple times, the polished consensus sequences produced during each round were aligned with MAFFT, after setting the gap penalty to 0. The final consensus was retrieved using EMBOSS cons based on the majority rule, namely including a base in the final consensus if it was included in at least 50% of the iterations. PCR primers were trimmed from both sides of the consensus sequence using Seqtk trimfq. As a final step, the consensus sequences were aligned using Blast v2.2.28+ against the NCBI nt database, which was downloaded locally. Seeds for subsampling reads in the three iterations reported in the results were 1, 3 and 5 in the draft consensus step, and 2, 4 and 6 for the polishing step, respectively. The accuracy of MinION consensus sequences was evaluated by aligning the *ONTrack* consensus sequence to the corresponding Sanger-derived reference sequence using Blast v2.2.28+ [17]. The accuracy of MinION reads was evaluated by aligning them to the corresponding Sanger reference sequence using Minimap2 and running Samtools stats on the generated *bam* file.

All scripts were run within an Oracle Virtualbox v5.1.26 virtual machine emulating an Ubuntu operating system on a Windows laptop without using any internet connection, and are available at https://github.com/MaestSi/ONTrack.git. MinION-based consensus sequences and Sanger consensus sequences are available as Supplementary Materials.

Sanger, MinION and consensus sequences are available at GenBank under the BioProject PRJNA539982.

## 3. Results

### 3.1 COI barcode sequencing

To perform barcode sequencing in the field, the portable genomics laboratory we previously described [1] was optimized further to include equipment and reagents with greater stability and better performance in tropical environments (up to 35°C and 90% humidity) after transport on standard domestic and international flights. Currently, the laboratory comprises seven portable devices that can be fitted in one standard luggage item with dimensions of 55×45×20 cm (Figure 1).

After collecting two snails and five insects during a workshop held by Taxon Expeditions (https://taxonexpeditions.com/) at the Ulu Temburong National Park (Borneo, Brunei) in October 2018, we dissected the tissue and extracted DNA. PCR products obtained by amplifying ∼710 bp of the COI gene were sequenced in the field using the MinION device with R9.4 sequencing chemistry. The MinION flow cell showed 995 active pores during the pre-run quality control (starting from 1005 on delivery by the manufacturer) and produced 600,000 reads in 3.5 h. Raw fast5 reads were basecalled, demultiplexed and trimmed offline, resulting in 9,000–77,000 reads per sample (Table 1). When we returned to Europe, the same genomic fragments were amplified and sequenced from the same DNA extracts using the Sanger method to evaluate the accuracy of the MinION-based barcoding pipeline.

**Table 1.**
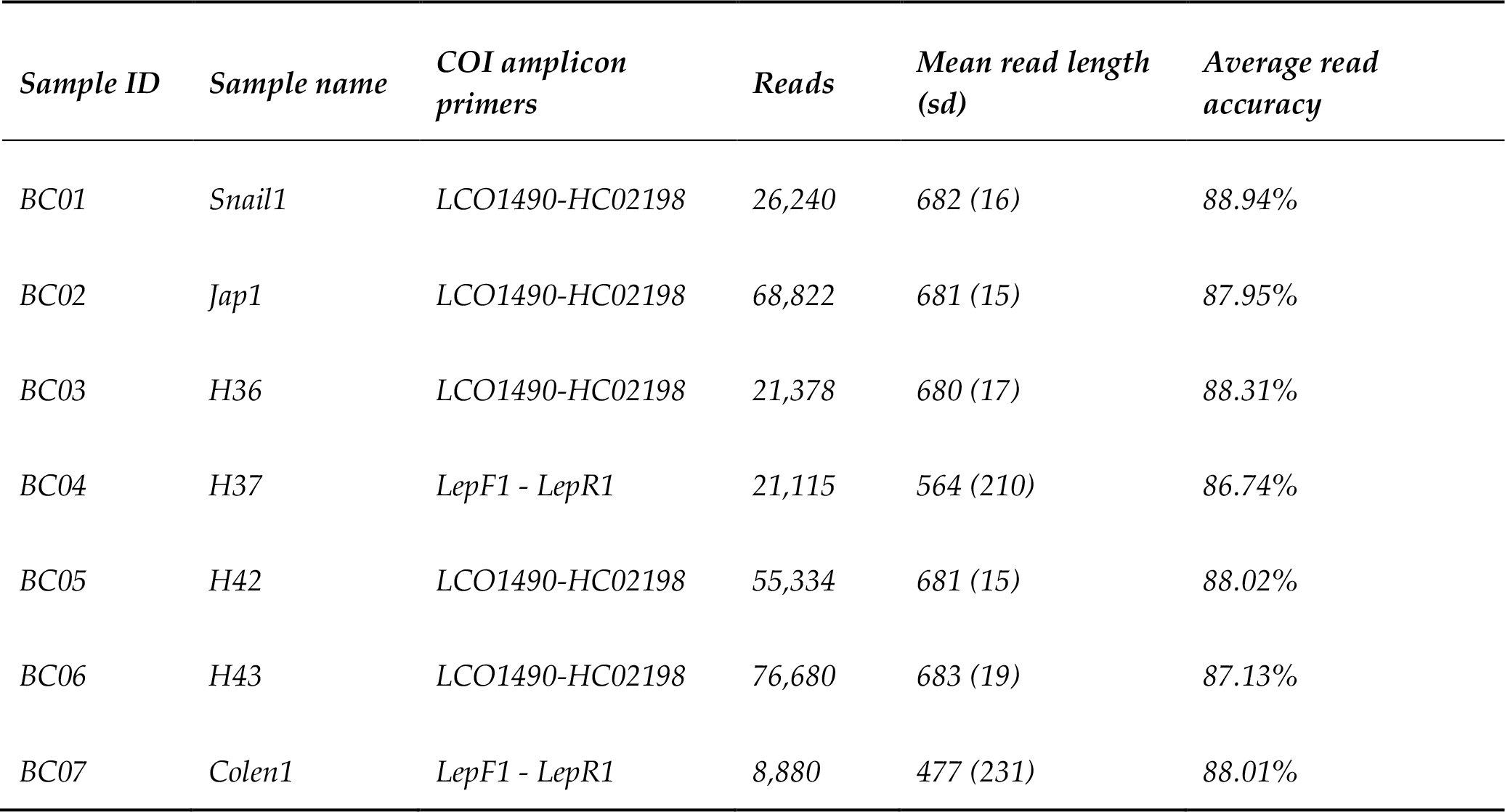
Sequencing statistics. For each sample, we show the COI primers used for PCR amplification, the number of sequenced reads, the mean and the standard deviation of read length in base pairs, and the average accuracy of MinION reads.

| *Sample ID* | *Sample name* | *COI amplicon primers* | *Reads* | *Mean read length (sd)* | *Average read accuracy* |
| --- | --- | --- | --- | --- | --- |
| BC01 | *Snail1* | LCO1490-HC02198 | 26,240 | 682 (16) | 88.94% |
| BC02 | *Jap1* | LCO1490-HC02198 | 68,822 | 681 (15) | 87.95% |
| BC03 | *H36* | LCO1490-HC02198 | 21,378 | 680 (17) | 88.31% |
| BC04 | *H37* | *LepF1 - LepR1* | 21,115 | 564 (210) | 86.74% |
| BC05 | *H42* | LCO1490-HC02198 | 55,334 | 681 (15) | 88.02% |
| BC06 | *H43* | LCO1490-HC02198 | 76,680 | 683 (19) | 87.13% |
| BC07 | *Colen1* | *LepF1 - LepR1* | 8,880 | 477 (231) | 88.01% |

### 3.2 Barcode analysis using the *ONTrack* pipeline

The MinION reads were processed using *ONTrack*, a barcoding pipeline that we developed using several samples collected over the last few years (Figure 2). The first step of the pipeline involved clustering the reads to remove non-specific PCR products and nuclear mitochondrial DNA segments (NUMTs), which can cause barcoding issues particularly when processing insect samples [18,19]. We then randomly sampled 200 of the filtered reads and aligned them to produce a draft consensus sequence. Starting from the draft consensus sequence, a polishing step was performed using another set of 200 randomly sampled reads.

**Figure 2:**
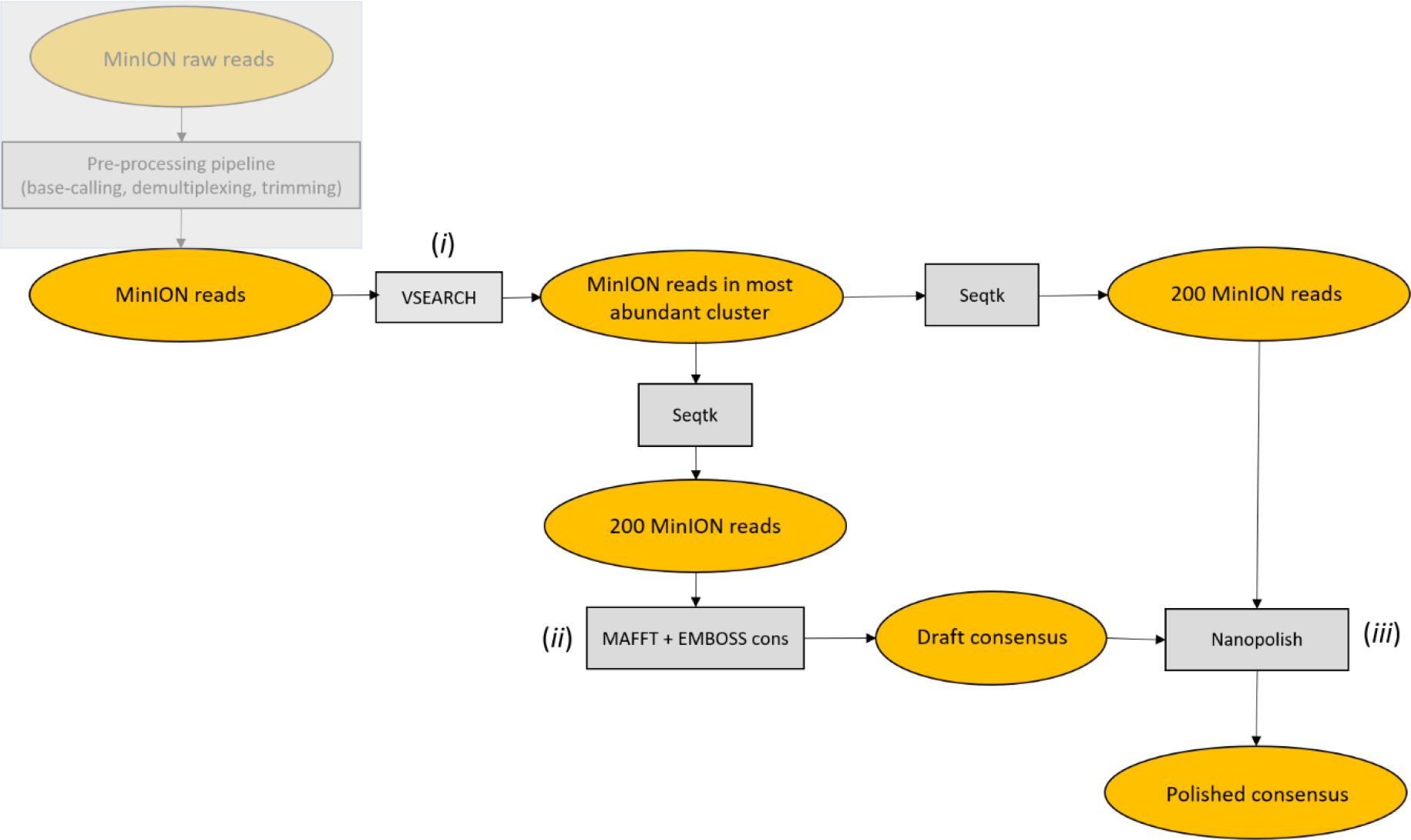
ONTrack pipeline flowchart. (i) MinION reads are clustered at 70% identity using VSEARCH and only reads in the most abundant cluster are retained for subsequent analysis. (ii) Next, 200 reads are then subsampled by Seqtk, aligned with MAFFT and a draft consensus is extracted with EMBOSS cons. (iii) The draft consensus sequence is then polished using Nanopolish, based on a second set of 200 randomly sampled reads.

Despite the errors characterizing MinION reads (Table 1), the barcodes reconstructed using the *ONTrack* pipeline had an average accuracy of 99.94% compared to the Sanger reference sequence. No consistent differences were observed between the two distinct types of COI amplicons we analyzed or the type of starting samples (Table 2).

**Table 2.** Accuracy of consensus sequences generated by the ONTrack pipeline. For each sample, we show the mean percentage accuracy of the consensus sequences obtained.

| *Sample ID* | *Consensus accuracy* |
| --- | --- |
| BC01 | 99.90% |
| BC02 | 100% |
| BC03 | 99.90% |
| BC04 | 100% |
| BC05 | 99.95% |
| BC06 | 99.89% |
| BC07 | 99.94% |

The generated consensus sequences were finally used as BLAST queries against the NCBI nt database, and the top hits for each sample were saved to a text file for operator analysis. Because the database was downloaded locally, the whole pipeline from sequencing to the generation of consensus sequences and the identification of BLAST top-hits could be completed without an internet connection, which was in any case unavailable in the field on our expedition.

We found that, when running the *ONTrack* pipeline three times for the same sample, the results differed slightly each time with an average accuracy ranging from 99.91% to 99.95%, depending on the read group subsampled in each analysis (Table 3). The pipeline was therefore run iteratively by aligning the consensus sequences generated during each round and extracting the ultimate consensus sequence. This slightly increased the accuracy of our barcoding pipeline, removing errors present in only one of the three iterations and thus achieving an average accuracy of 99.95%. The residual errors were only present in homopolymer runs of at least 6 nt, although some homopolymer runs of 7 nt were correctly reconstructed (Figure 3). The computational running time scaled linearly with the number of iterations, making it feasible to perform three iterations in a reasonable amount of time (∼30 min per sample) on a standard laptop.

**Table 3.** Accuracy of consensus sequences generated by combining three iterations of the ONTrack pipeline. For each sample, we show the number of properly reconstructed positions divided by the alignment length and (in parentheses) the percentage accuracy of the consensus sequences for each of the three iterations, the final consensus accuracy and the number of iterations supporting it.

| *Sample ID* | *Consensus accuracy read set 1* | *Consensus accuracy read set 2* | *Consensus accuracy read set 3* | *Final consensus accuracy* | *Iterations supporting the final consensus* |
| --- | --- | --- | --- | --- | --- |
| BC01 | 650/651 (99.85%) | 651/651 (100%) | 650/651 (99.85%) | 650/651 (99.85%) | 2/3 |
| BC02 | 656/656 (100%) | 657/657 (100%) | 657/657 (100%) | 657/657 (100%) | 3/3 |
| BC03 | 647/64 (100%) | 646/647 (99.85%) | 646/647 (99.85%) | 647/647 (100%) | 1/3 |
| BC04 | 606/606 (100%) | 606/606 (100%) | 606/606 (100%) | 606/606 (100%) | 3/3 |
| BC05 | 656/656 (100%) | 656/656 (100%) | 657/658 (99.85%) | 656/656 (100%) | 2/3 |
| BC06 | 576/576 (100%) | 575/576 (99.83%) | 574/575 (99.83%) | 575/576 (99.83%) | 2/3 |
| BC07 | 535/536 (99.81%) | 536/536 (100%) | 536/536 (100%) | 536/536 (100%) | 2/3 |

**Figure 3.**
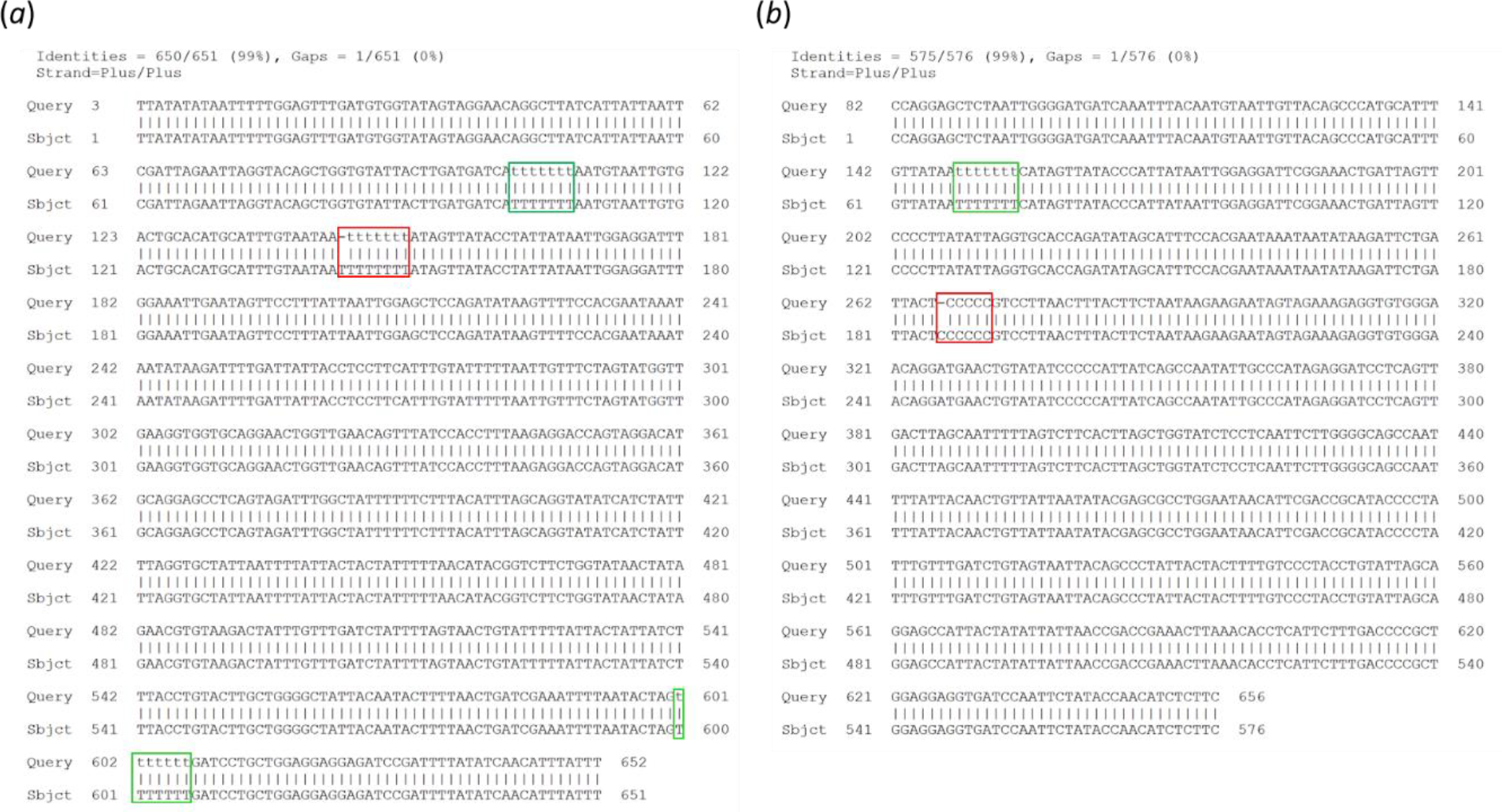
Analysis of residual errors in the ONTrack final consensus sequences. Alignment of the MinION consensus sequence (Query) to the Sanger sequence (Sbjct) is shown for samples BC01 (a) and BC06 (b). The residual errors, present in homopolymer runs of 6 and 8 nt, are highlighted in red. Properly reconstructed homopolymers of 7 nt are highlighted in green.

## 4. Discussion

We have described the implementation of a new workflow for barcoding in the field, from DNA extraction to the generation of consensus sequences. The selected protocols allowed the extraction of DNA from tiny snail-tissue biopsies and from whole beetles after cutting the abdomen to release soft tissues, as required to preserve the integrity of the specimens for detailed morphological evaluation. PCR products were successfully obtained despite the transport of our equipment in a standard Peli case and the storage of molecular biology reagents in local fridges and freezers powered for only 10 h per day. The MinION flow cells, which were not adversely affected by the transportation and storage conditions, retained most of their active pores and produced a good number of reads in a few hours. These results indicate that the molecular biology field laboratory workflow was robust, allowing us to barcode organisms at the collection site even under adverse environmental conditions (in this case a rainforest characterized by high temperatures and humidity).

On the software side, the new bioinformatics pipeline allowed us to analyze MinION reads using open-source and custom-developed scripts that run locally on a Linux Virtual Machine. The sequencing and data analysis could therefore be combined on a standard Windows laptop with a user-friendly interface. Most importantly, the improvements addressed some of the weaknesses of earlier pipelines, such as their dependence on sequence databases and Q-score calibration. The *ONTrack* pipeline works with as few as ∼500 reads per sample and achieves high accuracy when applied to MinION sequencing data obtained from COI barcode amplicons. Moreover, starting from processed MinION reads, the *ONTrack* pipeline returns consensus sequences in a few minutes, making it particularly suitable for work in the field.

The residual error rate in our consensus sequences never exceeded ∼0.2%. The proposed workflow can therefore be considered as a powerful tool for species identification given that most species pairs show sequence divergence exceeding 2% [7]. Further improvements may be achieved thanks to the software and chemistry enhancements regularly provided by ONT. A new flip-flop basecalling algorithm (https://github.com/nanoporetech/flappie) was recently implemented in the Guppy production basecaller and it should further reduce the error rate, albeit at the expense of basecalling time. A new sequencing chemistry (R10) will be released soon, increasing the accuracy especially in homopolymer runs and thus bringing on-site sequencing ever closer to the quality of Sanger analysis.

Sequencing and basecalling currently remain the most time-consuming steps in the pipeline, but both the hardware and software solutions provided by ONT are likely to become much more agile in the near future. Indeed, ONT recently released MinIT, a rapid analysis and device-control accessory for nanopore sequencing that connects to the MinION sequencer and performs GPU-accelerated and real-time basecalling. Moreover, the Medaka tool (https://github.com/nanoporetech/medaka) is expected to create polished consensus sequences faster than Nanopolish because it starts from basecalled data rather than raw signals. Finally, new MinION flow cells (Flongle) were recently made available and these are suitable for experiments that do not require a massive throughput, thus substantially reducing sequencing costs for small datasets. Because the *ONTrack* pipeline provides high-quality results with as few as ∼500 reads per sample (0.35 Mbp), multiple samples could be multiplexed in a single run and still fit Flongle specifications (1 Gbp) further reducing the cost. Considering a multiplex of 12 samples in a Flongle run, currently the maximum supported by standard ONT kits, we estimated a cost of about 30 € per sample to generate a barcode sequence with the workflow described herein. This is not far from the costs of standard Sanger sequencing (∼15 € per sample when sequencing both strands, without considering the extra shipment costs). Remarkably, the entire portable genomics laboratory described in this article can be acquired with a modest budget of 6000 €, compared to ∼80,000 € for a Sanger sequencer (ABI capillary). Dedicated, expert personnel are required to run the latter instrument, whereas the MinION sequencer is very simple and requires no special training. An additional significant advantage is that, unlike other sequencing technologies, the real-time MinION device does not require the number of sequenced reads to be set before the experiment begins. Therefore, the sequencing run can be stopped at any time when the necessary number of reads has been generated, achieving further cost and time savings.

## Author Contributions

Conceptualization, M.D. and M.S.; methodology, S.M., E.C., M.M., M.R. and M.D.; software, S.M.; validation, S.M. and E.C.; formal analysis, S.M. and E.C.; investigation, E.C., M.R., M.P., H.F., M.A., I.N., L.M., M.S., F.S., J.G.; writing—original draft preparation, S.M., M.R. and M.D.; writing—review and editing, M.R. and M.D.; visualization, S.M.; supervision, M.R. and M.D.; project administration, M.R. and M.D.; funding acquisition, M.D., M.S. and I.N.

## Acknowledgments

We gratefully acknowledge the Ulu Temburong National Park (Brunei, Borneo) for permission to conduct research in the field; export of biological materials was done under permit BioRIC/HoB/TAD/51 from the Ministry of Primary Resources and Tourism, Brunei Darussalam. We thank Davide Canevazzi for the support in bioinformatic analysis.

